# Coronary Artery Disease Risk Variant rs6903956 Links to Endothelial Dysfunction via *PHACTR1* Regulation

**DOI:** 10.1101/2025.05.11.653298

**Authors:** Kai Yi Tay, Hannah Su-Ann Wee, Nhi Nguyen, Matias Ilmari Autio, Vanessa Kristina Wazny, Khang Leng Lee, Darwin Tay, Yuting Wang, Xu Gao, Chew Kiat Heng, Mark Yan Yee Chan, Roger Sik Yin Foo, Jimmy Lee, Marie Loh, Christine Cheung

## Abstract

Ischemic heart disease, particularly coronary artery disease (CAD), remain leading causes of mortality worldwide. The single nucleotide polymorphism rs6903956 on chromosome 6p24.1 has been identified as a susceptibility locus for CAD in East Asian populations through genome-wide association studies. However, its functional role has not been fully elucidated. This study investigates the mechanistic basis of rs6903956 and its contribution to CAD pathogenesis, focusing on endothelial cell dysfunction. We first conducted cohort studies, revealing an association between the rs6903956 ‘A’ risk allele and blood pressure phenotypes, along with impaired endothelial responsiveness indicated by reduced flow-mediated dilation. Single-base editing of induced pluripotent stem cell-derived endothelial cells obtained from patients with CAD and expression quantitative trait loci analysis highlighted a cis-acting impact of the ‘A’ allele on *PHACTR1* and *EDN1* expression, suggesting allelespecific regulatory effects. Using *in silico* modeling by AlphaFold 3 platform, the ‘A’ allele exhibited enhanced binding affinity for HOXA4 and MEIS1 transcription factors, forming a stable ternary complex that promoted transcriptional activation of *PHACTR1*. Functional assays demonstrated the enhancer role of rs6903956 ‘A’ in *PHACTR1* promoter activity, supporting its locus-specific regulatory function in endothelial cells. Under pathological flow conditions, endothelial cells harboring the ‘A’ allele display elevated ICAM-1 expression and increased monocyte adhesion compared to the ‘G’ allele, indicating allele-specific endothelial inflammatory activation. These findings propose a model in which rs6903956 influences *PHACTR1* expression via HOX-MEIS cooperative binding, thereby modulating endothelial function and contributing to CAD susceptibility. This study provides mechanistic insights into the role of rs6903956 in endothelial dysfunction and CAD, informing potential therapeutic targets arising from genetic determinants in cardiovascular pathogenesis.

## Introduction

Coronary artery disease (CAD) is the leading cause of morbidity and mortality worldwide. Genome-wide association studies (GWAS) have identified over 300 genetic loci associated with CAD susceptibility^1^. Among these, the genetic variant rs6903956, located in the first intron of the *androgen-dependent tissue factor pathway inhibitor regulating protein* (*ADTRP*) gene on chromosome 6p24.1, has been linked to an increased risk of CAD. The minor allele ’A’ of rs6903956 is consistently associated with CAD in multiple studies conducted across East Asian populations^2–6^.

ADTRP regulates tissue factor pathway inhibitor expression in endothelial cells through an androgen-dependent mechanism^7^. Its knockdown promotes monocyte adhesion, LDL accumulation, and hallmarks of atherosclerosis, such as leaky vessels and inflammation^8,9^. Lower plasma ADTRP levels in CAD patients may suggest its cardio-protective role^10^. However, the functional role of rs6903956 and its impact on *ADTRP* remain poorly understood. An earlier study demonstrated that GATA2 binding to a 519 bp region containing the rs6903956 non-risk allele ‘G’ enhances *ADTRP* expression in HeLa cells, suggesting that rs6903956 may function as an enhancer^8^. On the other hand, rs6903956 may regulate the expression of a distant gene, *CXCL12*, on chromosome 10q11.21 through a trans-acting mechanism, potentially linked to endothelial activation pathways^11^. In endothelial cells, chromosomes 6p24.1 and 10q11.21 are spatially organized, bringing 6p24.1 near a weak promoter of *CXCL12*. *CXCL12*, a chemokine, plays a key role in atherosclerosis by interacting with the classical CXCR4 and atypical ACKR3 (CXCR7) receptors^12^. While GWAS-identified risk variants in intronic regions could interact with cell-type-specific enhancer elements to regulate the expression of distal genes^13^, it is crucial to investigate the cis-regulatory effects of rs6903956 and other genes within the 6p24.1 locus. This requires the application of advanced genome-editing technologies and specialized cellular models, such as arterial endothelial cells.

In this study, we highlight the role of rs6903956 as a potential modulator of blood pressure and endothelial function, supported by analyzes of the UK Biobank’s Phenome-wide Association Study, the Multi-Ethnic Study of Atherosclerosis in the USA, and replicated in a Han Chinese cohort from Guangxi China. Experimentally, we used single-base editing to convert the rs6903956 AA risk genotype in CAD patient-derived induced pluripotent stem cells (iPSCs) to the GG non-risk genotype and differentiated them into arterial endothelial cells. The AA genotype significantly increases *endothelin-1* (*EDN1*) and *phosphatase and actin regulator 1* (*PHACTR1*) transcript levels in endothelial cells, revealing its regulatory effect on neighboring cisgenes. *In silico* analyses and AlphaFold predictions show that the rs6903956 ‘A’ risk allele binds to transcription factors from the HOX and TALE families, with the HOXA4-MEIS1 complex specifically binding to the ‘A’ allele. Mechanistically, in endothelial cells with the ‘A’ risk alleles, HOXA4 knockdown reduced *PHACTR1* expression. Dual luciferase assays confirmed the enhancer role of the rs6903956 region, showing increased *PHACTR1* promoter activity with HOXA4 and MEIS1. Furthermore, endothelial cells with the AA risk genotype showed higher ICAM-1 expression and increased monocyte adhesion under disturbed flow conditions compared to those with the GG non-risk genotype. In summary, we link the allelespecific functional effects of rs6903956 to endothelial inflammatory activation through HOXA4-MEIS1-mediated regulation of *PHACTR1* expression.

## Methods

### Study approvals and subject enrolment

This study was approved by the Local Ethics Committee of Nanyang Technological University Singapore Institutional Review Board (IRB18/09/02 and IRB-2020-09-011), and the National Healthcare Group (DSRB: 2013/00937), Singapore. This research complies with the Helsinki Declaration. Written informed consent was obtained from each participant after the nature and possible consequences of the studies have been explained. We leveraged on the ‘Singapore Coronary Artery Disease Genetics Study’ where CAD patients from the National University Hospital angiography center had been genotyped for rs6903956 alleles. CAD patients were diagnosed as non-ST-elevation myocardial infarction (NSTEMI) by angiography.

### MESA study population and phenotypic characterization

The Multi-Ethnic Study of Atherosclerosis (dbGAP study accession phs000420.v6.p3) is an observational cohort study designed to examine the determinants of subclinical cardiovascular diseases in adults aged 45–84 years. It involves a diverse cohort of 6,814 asymptomatic men and women aged 45-84. 38 percent of the recruited participants are white, 28 percent African American, 22 percent Hispanic and 12 percent Asian, predominantly of Chinese descent. Participants free of clinical cardiovascular diseases at baseline were recruited from 6 field centers (New York, New York; Baltimore City and County, Maryland; Forsyth County, North Carolina; St. Paul, Minnesota; Chicago, Illinois; and Los Angeles County, California) between 2000 and 2002. More information about the objectives and design of MESA can be obtained from Bild et al., 2022^14^.

Brachial artery FMD was ascertained *via* ultrasound in MESA participants at baseline^15^. A linear-array multifrequency transducer operating at 9 MHz (GE Logiq 700 Device) was used to acquire images of the right brachial artery. After baseline images were obtained, the standard blood pressure cuff around the right arm was inflated to 50mm Hg above the participant’s systolic blood pressure for 5 minutes. Digitized images of the right brachial artery were captured continuously for 30 seconds before cuff inflation and for 2 minutes beginning immediately before cuff deflation to document the maximum vasodilator response. Change in FMD was calculated in 2 ways: absolute FMD (mm FMD), defined as maximum diameter— baseline diameter, and relative FMD (%FMD), calculated as ((maximum diameter − baseline diameter)/baseline diameter) × 100%. We reported associations with both FMD (mm) and % FMD.

### Retrieval of individual level genotypic information from MESA cohort and statistical analysis

Genotypic data were obtained from the database of Genotypes and Phenotypes (dbGaP) as the MESA SNP Health Association Resource (SHARe) project (study accession phs00209). Briefly, genotype calls were downloaded in matrix format and extracted from dbGAP using SRA toolkit. Affymetrix gene annotation files were used to map for rs6903956 and individual genotypes extracted using PLINK^16^.

Univariate analysis of covariance (adjusted for age and gender) was performed to evaluate the association between rs6903956 genotypes with FMD (mm) and %FMD. Analysis was done using SPSS statistics.

### Participants of Guangxi study and statistical analysis

The Guangxi Eco-Environmental Health and Aging Study (GEHAS) is an ongoing population-based cohort initiated between December 2018 and December 2022 in Guilin, Guangxi Zhuang Autonomous Region, China. The cohort targets middle-aged and older adults, with inclusion criteria as follows: (1) aged 45 years or older; (2) permanent residency in the region for at least three years; and (3) the ability to complete baseline and follow-up assessments. At enrollment, all participants underwent standardized evaluations, including structured interviews, physical examinations, and biospecimen collection.

The first follow-up survey was conducted between 2023 and 2024. A subset of 1,027 participants was randomly selected for genome-wide genotyping using the Illumina Asian Screening Array (ASA), which comprises approximately 700,000 markers and offers enhanced coverage of low-frequency variants relevant to East Asian populations. Genotyping data were processed and quality-controlled using PLINK (v1.9). Following standard quality control procedures, the SNP rs6903956 was extracted for downstream association analyses.

We assessed the association between the SNP rs6903956 and several cardiovascular-related physiological indicators using general linear regression models. The SNP rs6903956 was coded additively according to the number of “A” alleles (i.e., 0, 1, or 2). The outcome variables included systolic blood pressure (SBP), diastolic blood pressure (DBP), triglycerides (TG), low-density lipoprotein cholesterol (LDL-C), high-density lipoprotein cholesterol (HDL-C), hemoglobin A1c (HbA1c), and fasting blood glucose (FBG). All models were adjusted for age and sex.

Analyses were performed among 1,027 participants from the GEHAS cohort. Statistical significance was defined as a two-sided p-value <0.05.

### Characterization of induced pluripotent stem cells and arterial endothelial differentiation

CAD patient iPSCs were grown on Matrigel-coated plates (Corning, catalog no. 354230) in mTeSR1 medium. Cells were passaged every 4 to 5 days using ReLeSR (StemCell Technologies, catalog no. 05872). We performed characterization of our iPSCs by immunofluorescence, karyotyping and teratoma formation assay^11^. We followed our previously established differentiation protocols for arterial endothelial cells by Ang et al., 2022^17^.

### RNA extraction and quantitative RT-PCR

Total RNA was isolated using RNeasy Plus Mini kit (Qiagen, catalog no. 74134) as per the manufacturer’s protocol, subsequently used to generate cDNA with LunaScript RT SuperMix Kit (New England Biolabs, catalog no. E3010S). Real-time PCR was performed using SYBR green gene expression assays (New England Biolabs, catalog no. M3003S) on a QuantStudio 6 instrument (Applied Biosystems). Gene expressions were normalized to endogenous GAPDH housekeeping gene.

### Plasmid construction

All restriction enzymes were purchased from NEB, unless specifically stated. PCR reactions were conducted using Q5® Hot Start High-Fidelity 2X Master Mix (NEB, M0494L). Ligations were conducted using isothermal assembly with NEBuilder® HiFi DNA Assembly Master Mix (NEB, E2621L). All primers & oligos were ordered from Integrated DNA Technologies, Singapore. The pMIA20 plasmid used in this manuscript will be made available via Addgene. Primers used for fragment amplification are listed in Supplementary Table S5.

To build pMIA20 we first digested a pMIA3 (Addgene #109399)^18^ with Esp3I and ligated annealed oligos (Supplementary Table S5) to change the restriction cut sites at the gRNA ligation site into AarI-PaqCI. The U6-gRNA-scaffold expression cassette was amplified from the AarI-PaqCI modified pMIA3 plasmid. We digested pCMV_ABEmax_P2A_GFP (Addgene #112101, a kind gift from David Liu)^19^ with MluI and ligated the AarI-PaqCI-gRNA expression cassette, resulting in pMIA20 plasmid. gRNAs to target the SNP of interest were annealed together and cloned into pMIA20 digested with AarI (ThermoFisher Scientific, ER1581). The completed plasmid was used for base editing in the iPSC lines.

### Single base editing on CAD iPSCs

A candidate gRNA was designed to match the 5’-to-3’ DNA sequence containing rs6903956, located directly before an NGG PAM site: 5’-CATAGATTATTACTTAAGGT-3’. This gRNA duplex oligo was subcloned and ligated into the pMIA20 plasmid using T4 DNA ligase (New England Biolabs, catalog no. M0202S). 1.5x10^6^ cells were pre-treated with 10 μM ROCKi (StemCell Technologies, catalog no. 72302) for an hour. Single cell suspension was prepared using accutase (StemCell Technologies, catalog no. 07922) and resuspended in 100ul of nucleofection solution and 10ug of pMIA3 plasmid containing the selected gRNA pairs. Cells were nucleofected using Amaxa4D nucleofector (Lonza, catalog no. AAF-1002B) and P3 primary kit (Lonza, catalog no. V4XP-3024) as per manufacturer’s instructions and plated out on matrigel coated 6-well plate using mTeSR with CloneR supplement. After 48 hours, cells were again pre-treated with ROCKi and a single cell suspension was prepared. Fluorescence activated cell sorting (FACS) was performed to enrich targeted cells. RFP+ cells were sorted and plated onto 6-well plates containing mTeSR with CloneR and P/S. Colony formation was apparent from the individually sorted iPSCs 8 days after FACS. 24 single colonies then were manually picked into 12-well plates. Colonies were amplified and split with RelesR onto 6-well plates. Remaining cells were used for genomic DNA extraction and genotyping. Clones with successful CRISPR targeting were expanded.

### ChIP-Seq analysis

Histone marks in the 6p24 locus were queried from Roadmap epigenomics database by the ENCODE project. We downloaded the CTCF ChIP-Seq datasets from the ENCODE portal^20^ (https://www.encodeproject.org/) with the following identifiers: ENCFF047DPU, ENCFF105ZMP, ENCFF334OZC. Endothelial H3K4me3 and H3K27Ac ChIP-Seq datasets were downloaded from GEO under the accession number GSE131953^21^.

### TaqMan SNP Genotyping for rs6903956 base editing

Genomic DNA samples were extracted by DNeasy Blood and Tissue Kits (Qiagen, catalog no. 69506). TaqMan SNP genotyping assay was performed using predesigned TaqMan primers (Thermofisher, catalog no. 4351379) on a QuantStudio 6 instrument (Applied Biosystems). 20ng/μl of purified DNA sample was loaded into a 20ul reaction containing TaqMan Genotyping Master Mix (Thermofisher, catalog no. 4371353) on a 96 well plate format. The rs6903956 genotype of each sample was determined by the QuantStudio Design and Analysis Software available with the qPCR machine.

### Transcription factor binding site prediction

DNA motif of ∼60 bp flanking rs6903956 were used as input for Tomtom meme suite to identify potential transcription factor binding to rs6903956 region. Only the transcription factor associations p <0.001 with E-value <10 were considered as statistically significant.

### AlphaFold 3

To analyze the structural context of rs6903956, a 101 bp sequence centred on the SNP (TTCCATCTCAAAAATAAATAAATAAATAAATAAATAAATAGTGCCATAG(A/G)TTATT ACTTAAGGTTGGTCCCCCAAGTGTTGAAGTGGGTGCCATACTATAA) was input into the AlphaFold 3 model^22^ via the AlphaFold Server with the seed set to 1. The model generated five predictions, pre-ranked based on overall complex confidence using the ranking_score metric.

SASA was used to define the interaction residues on the protein complex interacting with the 101 bp DNA molecule centered on rs6903956. Residues were classified as interface residues if at least 15 Å² of their surface area was buried upon interaction with the DNA molecule. The buried SASA for the entire interface was quantified, and chain-chain interactions were considered significant if they resulted in a buried SASA of at least 300 Å². This analysis helped identify and characterize the residues involved in DNA-protein interactions.

Residue interactions were analyzed based on spatial proximity. Residues were classified as potential contact residues if the distance between them was within 8 Å. The distance between chains and individual residues was investigated to identify and characterize interaction interfaces.

### RNA-Seq processing

RNAseq libraries were prepared using samples of whole blood (n=1,234) collected in PaxGene RNA tubes at enrolment. RNAseq libraries were prepared from at least 1mg of total RNA using NEBNext® Ultra™ II Directional RNA Library Prep (New England Biolabs, Inc.), with GLOBINClear (Thermo Fisher Scientific) for depletion of globin gene RNA and Ribosomal RNA (rRNA). The libraries were sequenced on a NovaSeq6000, using a paired-end run of 2 x 150bp. We aimed for at least 30M aligned reads per library (∼9Gb of data). Adapter and quality trimming were performed using TrimGalore ^23^, whereas SortMeRNA ^24^ was used for the removal of rRNA. Alignment to the (GRCh38) reference genome was conducted using STAR version 2.7.9a ^25^, followed by quantification of reads with RSEM version 1.3.3 ^26^ ; which identified a total of 60,708 genes. Gender mismatch check was performed by interrogating for anomaly across 5 genes – namely XIST, RPS4Y1, EIF1AY, DDX3Y, and KDM5D. A total of 6 samples had failed this check; resulting in a total of 1,228 samples for downstream analysis. Finally, the genes were normalized using the Trimmed Mean of the M-values (TMM) ^27^ approach.

### Whole genome sequencing (WGS) processing

Genotyping was carried out from a combination of low coverage (15x) sequencing and imputation as detailed in the Yew et al.^28^. In brief, paired end 151bp WGS was performed on the Illumina HiSeq X with an average sequencing depth of 15.8X per sample (n=2,400). The TopMed Imputation Server was used to impute autosomal SNPs to the TopMed (Version R2) reference panel using the EAGLE2+Minimac4 pre-phasing and imputation pipeline. Approximately 285 million autosomal SNPs were available following imputation. Post-imputation quality control excluded imputed SNPs with MAF <0.001 in at least one of the three main ancestral groups (Chinese, Indians and Malays), as well SNPs with imputation INFO score <0.30 and RUTH HWE test P <10-3. A total of 7,857,631 imputed autosomal SNPs formed the final dataset, with 7,150,557 SNPs remaining with a MAF filter of 0.01. Genotype dosage values were computed using bcftools.

### Expression quantitative trait loci (eQTL)

eQTL analysis was performed using R package MatrixEQTL ^29^ whereby gene expression was modelled as a regression model of genotype dosage and 10 covariates - namely age, sex, ethnicity, RIN (RNA integrity number) and the top 6 PEER (Probabilistic Estimation of Expression Residuals) factors ^30^.

### ChIP-Taqman qPCR

ChIP-qPCR was performed using High-Sensitivity ChIP Kit (Abcam, catalog no. ab185913). WT (AA), Unedited (AA) and edited (GG) arterial endothelial cells were harvested at a density of 5 x 10^5^ cells per reaction. Cells were washed with PBS and cross-linked with 1% formaldehyde for 10 minutes at room temperature, followed by quenching with 1.25 M glycine. Cross-linked cells were then lysed, and chromatin was extracted. Chromatin was sheared using a probe sonicator set to 25% power output, applying 3-4 pulses of 10-15 seconds each, with 30-40 seconds rest on ice between pulses. The efficiency of shearing was confirmed via gel electrophoresis, ensuring that the chromatin fragments were between 100-700 bp with a peak size of 300 bp.

Chromatin immunoprecipitation (ChIP) was performed using 2 µg of sheared chromatin per well. Antibodies against HOXA9 (Santa Cruz Biotechnology, catalog no. sc-81291), HOXA4 (Santa Cruz Biotechnology, catalog no. sc-515418), MEIS1/2/3 (Santa Cruz Biotechnology, catalog no. sc-101850), APE1 (Abcam, catalog no. ERP18378) and HMGA1 (Abcam, catalog no. EPR22421) were used for the ChIP assays. The positive control used was an antibody against RNA polymerase II. The negative control was non-immune IgG, which serves as a baseline to measure non-specific binding. The antibodies were bound to strip wells coated with a protein A/G mix that has a high affinity for IgG antibodies. Chromatin samples were then added to these wells along with ChIP buffer, enrichment enhancer, and blocker solution, and incubated to allow antibody binding to the target chromatin fragments.

After incubation, the reaction wells were washed four times with 200 µL of Wash Buffer to remove non-specifically bound chromatin. This was followed by a single wash with DNA Release Buffer. Cross-links between DNA and proteins were reversed by incubating with RNase A solution at 42°C for 30 minutes, followed by Proteinase K at 60°C for 45 minutes. The DNA was then purified using a spin column and eluted with DNA Elution Buffer.

For quantitative PCR (qPCR) analysis, input DNA (i.e., a fraction of the chromatin before immunoprecipitation) was used as a control to normalize the data. The purified DNA was quantified using Sybr Green qPCR primers for Allele-Specific qPCR^8^ as well as TaqMan qPCR primers (Supplementary Table S5) amplifying a 147bp region flanking rs6903956 A and G.

qPCR reactions were run with an initial activation step at 95°C for 7 minutes, followed by 40 cycles of 95°C for 10 seconds, 55°C for 10 seconds, and 72°C for 8 seconds. The fold enrichment of target sequences was calculated using the amplification efficiency of the ChIP sample relative to the input DNA.

### siRNA knockdown of HOXA4, HOXA9, MEIS1, MEIS2, PBX2

Cells were seeded at a density of 1.3 x 10^5^ cells per well in 24-well plates and incubated for 24 hours. SMARTPool siRNAs targeting HOXA4, HOXA9, MEIS1, MEIS2, and PBX2 were used (Dharmacon, catalog no. L-011693-00-0005, L-006337-00-0005, L-011726-00-0005, L-011330-00-0005, E-011746-00-0005).

siRNAs were resuspended by preparing a 20 µM stock solution. 5 nmol of siRNA was dissolved in 250 µL of 1x siRNA buffer (Dharmacon, catalog no. B-002000-UB-100). The solution was gently mixed and incubated on an orbital shaker for 30 minutes at room temperature. The concentration of siRNA was confirmed using spectrophotometry at 260 nm. Resuspended siRNAs were aliquoted and stored at - 20°C.

For transfection, to obtain a final concentration of 5nM siRNA, 0.5 µL/well of 5uM stock siRNA was diluted in 49.5 µL of Opti-MEM^TM^ reduced serum medium (Thermo Fisher Scientific, catalog no. 31985062). 1 µL/well of DharmaFECT 1 transfection reagent (Dharmacon, catalog no. T-2001-03) was used. The siRNA and transfection reagent mixtures were incubated separately for 5 minutes before combining and incubating for an additional 20 minutes at room temperature to form siRNA-lipid complexes. Transfection medium was prepared by mixing the siRNA-lipid complexes with EGM2 containing 10% HI FBS, resulting in a final siRNA concentration of 5 nM. The existing culture medium was removed from the cells, and the transfection medium was added. Cells were incubated for 72 hours. siRNA knockdown was confirmed and *PHACTR1* transcript levels quantified using qPCR.

### Dual luciferase assay

The 519 bp ADTRP regulatory region flanking rs6903956 (-789 bp to +724 bp) (chr6:11,774,037–11,774,555) from Luo et al. was cloned into the pGL3 Control Vector (Addgene, Plasmid #212937) using KpnI and XhoI, upstream of the SV40 promoter and firefly luciferase coding region. Additionally, the same 519 bp region was cloned into the pGL3 Basic Vector (Addgene, Plasmid #212936) using XhoI and HindIII. Subsequently, a 1781 bp regulatory region of PHACTR1 (-766 bp to +1015 bp) (chr6:12,716,001–12,717,781) was subcloned into the pGL3-Basic vector. Cloning was performed by GenScript Biotech (Piscataway, NJ, USA).

The expression plasmids pcDNA3.1(-)-HOXA4 and pcDNA3.1(-)-MEIS1 were also constructed by GenScript Biotech.

HUVECs were seeded at a density of 5000 cells per well in a 96-well plate. 24 hours after seeding, cells were transfected with 50 ng of the respective reporter construct and 0.5 ng of the pNL1.1 PGK[Nluc/PGK] vector control (Promega, Madison, WI, USA, catalog no. N1441) using 0.5 μL/well of DharmaFECT 1 transfection reagent (Dharmacon, catalog no. T-2001-03). Salmon sperm DNA was used as carrier DNA to normalize plasmid DNA concentrations. Cells were harvested 72 hours post-transfection, and firefly and NanoLuc® luciferase levels were measured using the Nano-Glo® Dual-Luciferase® Reporter Assay System (Promega, catalog no. N1531). Firefly/NanoLuc® activity was determined for each sample using a Synergy H1 microplate reader (BioTek) and measured in triplicates.

### Endothelial-monocyte interaction assay in perfusion cell culture

Inverted μ-Slide γ-shaped slides (ibidi GmbH, catalog no. 80126) were coated with rat tail collagen I and seeded with iPSC-derived endothelial cells (1.5 x 10^6^ cells/ml) in Endothelial Cell Growth Medium (Lonza, catalog no. CC-3162) supplemented with 10% heat-inactivated FBS (Life Technologies, catalog no. 10500064) and ROCK inhibitor (1:1000 dilution). Cells were cultured at 37°C and 5% CO2 for 24 hours. THP-1 monocytes were maintained in RPMI 1640 (Gibco, catalog no. 11875093) with 10% heat-inactivated FBS. THP-1 monocytes (1.5 x 10^6^ cells/ml) were labeled with LuminiCell Tracker 670 (Millipore, catalog no. SCT011) for 30 minutes at 37°C, and resuspended in serum-free RPMI medium before the assay.

Using the ibidi Pump System, shear stress of 12 dyn/cm^2^ was applied for 24 hours with red perfusion reservoir sets (ibidi GmbH, catalog no. 10962). THP-1 monocytes were added to achieve a final volume of 12-13 ml of EGM2 supplemented with 10% heat-inactivated FBS. Post-assay, cells were fixed in 4% PFA (Nacalai Tesque, catalog no. 09154-85), permeabilized with 0.1% Triton X-100 (Bio Basic Inc., catalog no. TB0198), and blocked with 1% BSA (Sigma-Aldrich, catalog no. A1595-50ML). Cells were incubated with primary antibodies (Supplementary Table S3) overnight at 4°C and with secondary antibodies for 1 hour at room temperature. DAPI staining was performed before imaging on a Nikon Eclipse TiE microscope with Metamorph software.

### Statistical analysis

Statistical analyses were performed using GraphPad Prism version 9. Data normality was assessed using the Shapiro-Wilk test. For normally distributed data, unpaired Student’s two-tailed t-tests were used for comparisons between two groups, and one-way ANOVA with post-hoc Tukey’s tests were applied for datasets with more than two groups. For non-normally distributed data, the Mann–Whitney U test or Kruskal-Wallis test was used as appropriate. A p-value < 0.05 was considered statistically significant. Results are presented as mean ± standard deviation (SD). Details of the statistical tests applied to specific experiments are provided in the figure legends.

## Results

### Population-based studies identify rs6903956 association with blood pressure phenotypes and endothelial dysfunction

We leveraged extensive phenotypic data from large-scale cohort studies to investigate the association of the genetic variant rs6903956 with cardiovascular phenotypes and related risk factors. Our analysis aimed to delineate tissue-specific effects of this variant. Firstly, we conducted analysis of a Phenome-wide Association Study (PheWAS) from the UK Biobank, a prospective cohort comprising approximately 500,000 individuals aged 40–69 from across the United Kingdom^31^. Our analysis focused on clinical diagnoses related to the circulatory system and identified significant associations between the rs6903956 variant and essential hypertension (*p* = 0.023) as well as general hypertension (*p* = 0.025) (**Supplemental Table 1**). No significant association was observed between rs6903956 and coronary atherosclerosis in our analysis. This lack of association may reflect the underrepresentation of Asian populations in the UK Biobank cohort, where the variant was originally identified as a risk locus for coronary artery disease in East Asian populations^2-6^. Building on the observed associations with hypertension, we further investigated traditional cardiometabolic risk factors in a cohort of 1,027 individuals from Guilin, Guangxi, China. Using linear regression analysis, we identified a significant association between the number of ’A’ alleles at rs6903956 and elevated diastolic blood pressure, with a regression coefficient of 2.31 (*p* = 0.045) after adjusting for gender and age (**Table 1**). These findings suggest potential allele-specific effects on blood pressure regulation.

**Table 1.** Associations Between Cardiometabolic Risk Factors and rs6903956 in a Population from Guilin, Guangxi, China. This table summarizes the associations of key cardiometabolic risk factors with rs6903956 in 1,027 individuals from the Guilin, Guangxi, China cohort. Effect sizes are presented with their respective standard errors (SE) and *p*-values, adjusted for gender and age. The analyzed outcomes include systolic blood pressure (SBP), diastolic blood pressure (DBP), triglycerides (TG), low-density lipoprotein cholesterol (LDLC), high-density lipoprotein cholesterol (HDLC), Hemoglobin A1C (HbA1C), and blood glucose levels (GLU).

| **Associations Between Cardiometabolic Risk Factors and rs6903956** |  |  |
| --- | --- | --- |
| **Risk factors** | **Effect size (SE)** | ***p*-value** |
| **LDLC** | **-0.007 (0.084)** | **0.929** |
| **TG** | **0.178 (0.125)** | **0.153** |
| **HDLC** | **-0.044 (0.041)** | **0.287** |
| **GLU** | **0.125 (0.167)** | **0.454** |
| **HbA1C** | **0.082 (0.105)** | **0.431** |
| **SBP** | **-0.093 (2.839)** | **0.974** |
| **DBP** | **2.310 (1.151)** | **0.045** |

Variations in blood pressure are often accompanied by vascular adaptations, including structural remodeling, endothelial dysfunction, and heightened vascular reactivity. To explore the vascular-specific impacts of the rs6903956 variant, we conducted a meta-analysis using data from the Multi-Ethnic Study of Atherosclerosis (MESA) cohort (dbGAP study accession: phs000420.v6.p3) ^14^, which investigates risk factors for subclinical cardiovascular diseases in a multi-ethnic population. The MESA cohort comprises 6,814 asymptomatic individuals from four ethnic groups: 38% White, 28% African American, 22% Hispanic, and 12% Asian. We focused on physiological measurements of vascular function and structure, analyzing associations of rs6903956 with arterial stiffness, carotid intima-media thickness, and flow-mediated vasodilation in 566 Asian American participants (**Table 2**). Flow-mediated vasodilation (FMD), expressed as a percentage (%), is a widely recognized predictor of cardiovascular events and an established method for assessing endothelial function by evaluating vascular responsiveness to changes in blood flow^15,32^. In our analysis, the rs6903956 A risk allele was significantly associated with reduced FMD (%) compared to the G allele (A allele: 3.959% [95% CI: 3.313–4.605] vs. G allele: 4.812% [95% CI: 4.585–5.038], p = 0.015, dominant model). Similarly, the A allele was significantly associated with decreased absolute FMD (mm) (A allele: 0.163 mm [95% CI: 0.138–0.187] vs. G allele: 0.197 mm [95% CI: 0.188–0.205], p = 0.010, dominant model). The observed reductions in both % FMD and absolute FMD (mm) in individuals with either dominant or additive model of A allele suggest impaired endothelial responsiveness (**Table 2**), linking rs6903956 to endothelial dysfunction. For arterial stiffness metrics—including aortic distensibility, large artery elasticity index, and small artery elasticity index—we found no significant associations under either additive or dominant models for the risk allele ’A.’ Similarly, no significant associations were observed between rs6903956 and carotid intima-media thickness. These findings suggest that rs6903956 does not significantly influence vascular stiffness or structural biomarkers in this population.

**Table 2.** Associations Between rs6903956 Genotypes and Vascular Function Parameters in the MESA Cohort. This table presents the associations of rs6903956 genotypes with physiological measurements of vascular function in 566 Chinese American participants from the Multi-Ethnic Study of Atherosclerosis (MESA) cohort (dbGAP accession: phs000420.v6.p3). Vascular function metrics analyzed include arterial stiffness parameters (aortic distensibility, large artery elasticity index, small artery elasticity index), carotid intima-media thickness (CIMT), and flow-mediated vasodilation (%FMD and mm FMD). Associations were assessed under dominant and additive genetic models using univariate analyses of covariance, adjusting for age and gender. Results are expressed as standardized coefficients (Beta), t-values, and p-values. Genotype distributions are as follows: GG (n = 504), GA (n = 59), AA (n = 3).

| Physiological measurements of<br>vascular function | Genetic model | Standardized coefficients |  |  |
| --- | --- | --- | --- | --- |
|  |  | Beta | t-value | p-value |
| Aortic distensibility | Additive | 0.113 | 1.570 | 0.118 |
|  | Dominant | 0.113 | 1.570 | 0.118 |
| Large artery elasticity indices | Additive | 0.052 | 1.064 | 0.288 |
|  | Dominant | 0.039 | 0.804 | 0.422 |
| Small artery elasticity indices | Additive | -0.017 | -0.342 | 0.733 |
|  | Dominant | -0.017 | -0.351 | 0.726 |
| Carotid Intima-Media Thickness | Additive | 0.042 | 0.753 | 0.452 |
|  | Dominant | 0.048 | 0.870 | 0.385 |
| % Flow-mediated vasodilation | Additive | -0.092 | -2.072 | 0.039 |
|  | Dominant | -0.108 | -2.445 | 0.015 |
| Flow-mediated vasodilation (mm) | Additive | -0.091 | -2.099 | 0.036 |
|  | Dominant | -0.112 | -2.579 | 0.010 |

Taken together, the data from three independent cohorts highlight the intriguing physiological role of rs6903956 as a potential modulator of blood pressure and endothelial function. These findings underscore the importance of endothelial cells as a primary target for subsequent mechanistic investigations. Our next section will focus on elucidating the functional genetics of rs6903956 and unraveling the fundamental endothelial biology influenced by this variant.

### Cis-acting impact of rs6903956 on PHACTR1 and EDN1 expressions uncovered by single-base editing in patient endothelial cells

Given the endothelial-specific effects associated with rs6903956, we aimed to investigate its precise mechanism using a human-relevant endothelial cell model. To this end, we employed single-base editing to modify the genotype of iPSCs derived from CAD patients. iPSCs harboring the AA risk genotype were edited to convert the sequence to the GG non-risk genotype (**Fig. 1A**). The edited iPSCs were subsequently differentiated into endothelial cells, creating a robust isogenic model for dissecting the genetic basis of rs6903956 in endothelial dysfunction. To enable precise single-base editing at the rs6903956 locus, we developed an integrated system combining the D10A Cas9 nickase, a guide RNA (gRNA) cassette, and an adenosine base editor (ABEmax) within a single plasmid construct (pMIA20) (**Supplemental Fig. 1A, B**). This design offered significant advantages over traditional base-editing approaches, which typically require the co-transfection of multiple plasmids to deliver Cas9, gRNA, and the base editor ^33^. By consolidating all components into a single construct, we enhanced transfection efficiency and minimized cellular stress, providing a more streamlined and effective approach to base editing.

**Figure 1.**
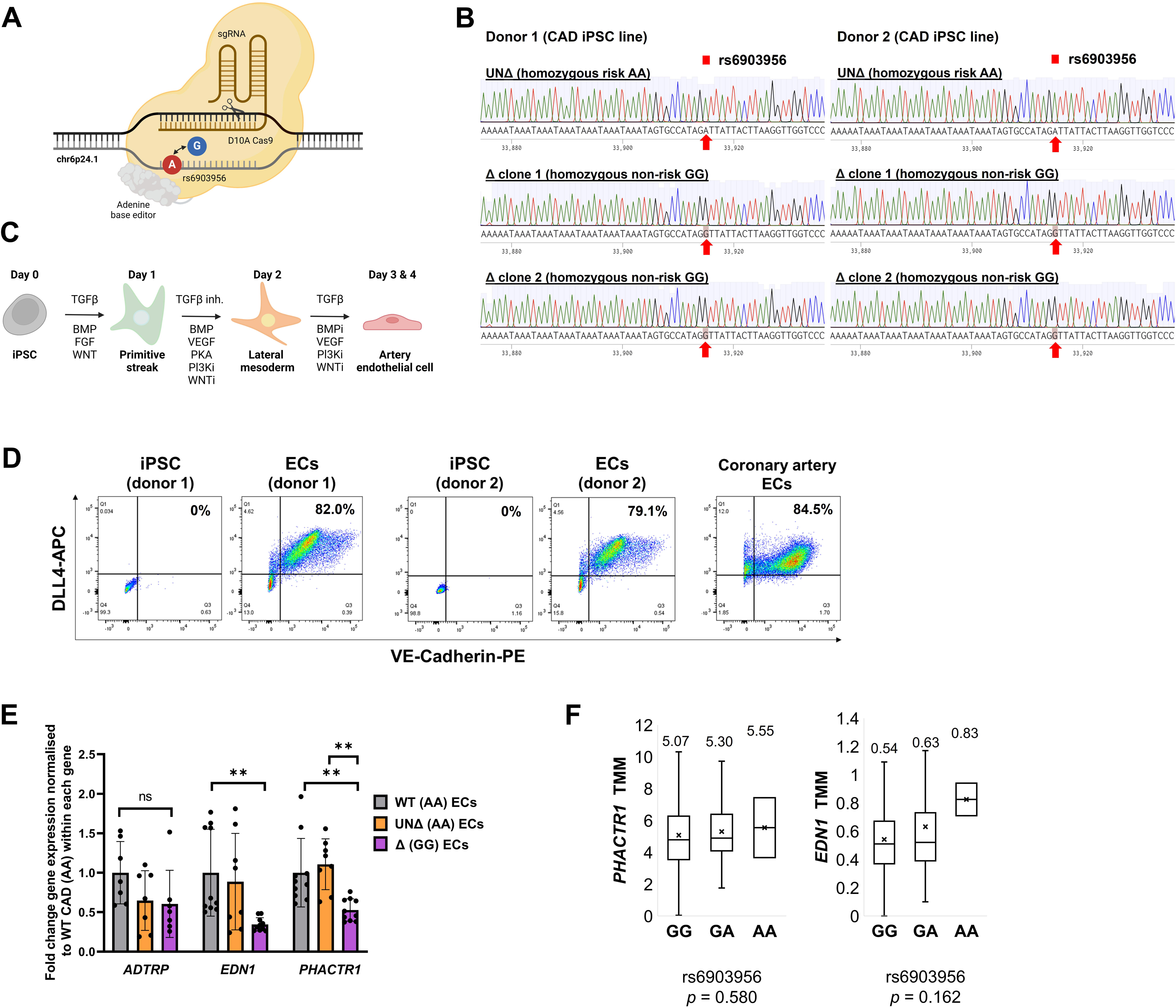
**Generation and Validation of iPSC-Derived Isogenic Cell Lines with rs6903956 Single-Base Edits from Coronary Artery Disease Patients.** (A) Schematic representation of single-base editing at rs6903956 using the pMIA20 plasmid containing D10A Cas9, an sgRNA targeting rs6903956, and an adenosine base editor. (B) Sequencing chromatograms confirming single-base edits at rs6903956 (hg38 chr6:11,714,312– 11,714,371). The rs6903956 variant has been highlighted by red arrows. Editing was performed on iPSCs derived from two CAD patients, generating two homozygous non-risk GG genotype clones (Δ clone 1 and Δ clone 2) per patient. An unedited control (UNΔ) carrying the homozygous risk AA genotype underwent pMIA20 nucleofection but did not result in successful editing. (C) Diagram of the 4-day differentiation protocol from iPSCs to endothelial cells, adapted from Ang et al., 2022. (D) Flow cytometry analysis validating endothelial cell identity. Differentiated endothelial cells were stained for the arterial marker DLL4 and endothelial marker CD144. Undifferentiated iPSCs served as negative controls, while human coronary artery endothelial cells served as positive controls. (E) Quantitative RT-PCR analysis of rs6903956 cis-regulated genes (*EDN1*, *PHACTR1*, and *ADTRP*) in WT (AA) - wild-type parental iPSCs with the AA risk genotype; Δ (GG) - base-edited iPSCs with the GG non-risk genotype; and UNΔ (AA) - unedited controls retaining the AA risk genotype. Data represent means ± S.D. from two donor cell lines from 2–3 differentiation batches, and two technical replicates per batch. Statistical significance was determined using within each gene, \*\**p* ≤ 0.01, Kruskal-Wallis test. (F) Expression quantitative trait locus (eQTL) analysis of *EDN1* and *PHACTR1* in the HELIOS cohort, Chinese participants adjusted for gender and age. Genotypes included GG (n = 749), GA (n = 99), and AA (n = 2). Gene expression levels were normalized using Trimmed Mean of M-values (TMM) and analyzed with an independent Kruskal-Wallis test.

Using this optimized system, we successfully performed base editing in iPSCs derived from two CAD patients (CAD donor 1 and CAD donor 2), converting the rs6903956 risk genotype (AA) to the non-risk genotype (GG) (**Fig. 1B**). To account for potential off-target effects and ensure experimental robustness, we generated control iPSCs by subjecting the same CAD-derived cells to the CRISPR-Cas9 nucleofection protocol without successful base editing. Initial screening was conducted using the TaqMan SNP genotyping assay, followed by Sanger sequencing to validate the precise base substitution and confirm the absence of unintended insertions or deletions (indels) (**Supplemental Fig. 1C**). The editing success rate ranged from 30–35% across both donors. To ensure consistency and suitability for downstream analyses, we selected two successfully edited clones and one unedited clone per donor. Selection criteria included the clarity of the Sanger sequencing results, maintenance of normal iPSC morphology, and absence of spontaneous differentiation. This approach facilitated the development of an isogenic CAD iPSC model, comprising three groups: WT (AA), representing the wild-type parental iPSCs with the AA risk genotype; Δ (GG), denoting successfully base-edited iPSCs with the GG non-risk genotype; and UNΔ (AA), serving as unedited controls retaining the AA risk genotype. We differentiated the isogenic iPSCs into endothelial cells (ECs) using an established protocol^17^ (**Fig. 1C**). An average of 80% of the differentiated ECs consistently coexpressed the endothelial marker VE-Cadherin and the arterial marker DLL4, as confirmed by flow cytometry (**Fig. 1D**). To ensure high-quality populations for downstream applications, cells were further purified through CD144+ (VE-Cadherin+) MACS sorting. The resulting endothelial cells were designated as WT (AA) ECs, UNΔ (AA) ECs, and Δ (GG) ECs, corresponding to their respective iPSC origins.

Since SNPs often regulate genes within the same topologically associating domain ^34^, we hypothesized that the rs6903956 A-to-G / G-to-A??? substitution might modulate the expression of cis-genes within the same domain. Hi-C contact maps, which reveal 3D chromatin architecture, indicate that rs6903956 resides within a topologically associating domain encompassing the genes *ADTRP*, *EDN1*, and *PHACTR1* ^11^. To investigate potential cis-regulatory effects of rs6903956, we analyzed the expression of these genes in WT (AA), UNΔ (AA), and Δ (GG) endothelial cells (**Fig. 1E**). While *ADTRP*, located within the first intron of rs6903956, has been associated with reduced expression due to the presence of ‘A’ risk allele in leukocytes^6^, no significant differences in *ADTRP* expression were observed among the three endothelial cell groups. This suggests that the regulatory effect of rs6903956 on *ADTRP* is likely cell-type specific and may not extend to endothelial cells. In contrast, we observed significant downregulation of *PHACTR1* and *EDN1* expression in Δ (GG) ECs compared to WT (AA) ECs, with *PHACTR1* expression also showing a significant difference between Δ (GG) ECs and UNΔ (AA) ECs. These findings suggest that *PHACTR1* could be a key target of rs6903956 in endothelial cells, potentially acting as a causal gene. Furthermore, the observed decrease in *EDN1* expression in Δ (GG) ECs may represent a downstream effect, as *PHACTR1* overexpression has been shown to upregulate *EDN1*^35^.

The *PHACTR1* locus at 6p24.1 has been identified as a risk locus for five vascular diseases, with the lead SNP rs9349379 associated with a putative enhancer for *EDN1*^36^. However, SNPs in *EDN1* have not been implicated in large-scale GWAS for CAD. In contrast, multiple studies have linked *PHACTR1* polymorphisms to blood pressure traits, including hypertension, systolic blood pressure, and pulse pressure^37–40^. Consistent with these findings, our analysis of the Guilin cohort identified a significant association between the number of ’A’ alleles at rs6903956 and elevated diastolic blood pressure (**Table 1**), suggesting that rs6903956 may regulate *PHACTR1* expression to influence blood pressure control. Importantly, genotyping of the cell lines used in our study confirmed the absence most of the *PHACTR1* risk alleles (**Supplemental Table 2**), ensuring that the observed effects could be attributable to rs6903956. Notably, lead SNPs on PHACTR1 in this region have also been linked to blood pressure-related phenotypes^37–40^.

We hypothesized that carriers of the ‘A’ risk allele would exhibit increased expression of *EDN1* and *PHACTR1*. To test this, we conducted an expression quantitative trait loci (eQTL) analysis using data from the Health for Life in Singapore (HELIOS) cohort, a longitudinal study of 10,004 individuals of Chinese, Indian, and Malay descent. In a subset of 850 Chinese participants, we analyzed gene expression profiles of peripheral blood mononuclear cells (PBMCs). After adjusting for age and gender, we observed a potential additive effect of the rs6903956 A risk allele on *EDN1* and *PHACTR1* expression levels (**Fig. 1F**). However, this trend did not reach statistical significance, possibly due to the limited number of ‘AA’ genotypes in the cohort or the cell-type specificity of rs6903956’s regulatory effects, which might obscure differences in PBMCs. Nonetheless, the observed trend of higher normalized *EDN1* and *PHACTR1* expression in ‘A’ allele carriers aligns with findings from our endothelial cell models, suggesting consistency across different biological contexts.

### In silico insights into rs6903956 ‘A’ allele-mediated cooperative binding of HOXA4 and MEIS1 in transcriptional regulation

The findings so far prompted us to explore the mechanisms through which rs6903956 might regulate *EDN1* and *PHACTR1* expression, given its location 512,763 kb from *EDN1* promoter and 941,651 kb from *PHACTR1* promoter. Variants such as rs6903956 may influence long-range gene regulation by altering transcription factor (TF) binding sites, thereby disrupting, creating, or modifying the affinity of these sites^41^. To investigate whether rs6903956 has a regulatory role, we assessed its potential impact on TF binding and cooperative enhancer activity. DNase hypersensitivity regions (DHSs), which mark active TF binding sites, provide critical insights into regulatory variants^42^. Using data from the ENCODE Project (GEO accession: GSM1014528), DNase I footprinting analysis in human umbilical vein endothelial cells (HUVECs) revealed that rs6903956 resides within a DHS (**Fig. 2A**). This suggests that rs6903956 may play a functional role in TF binding.

**Figure 2.**
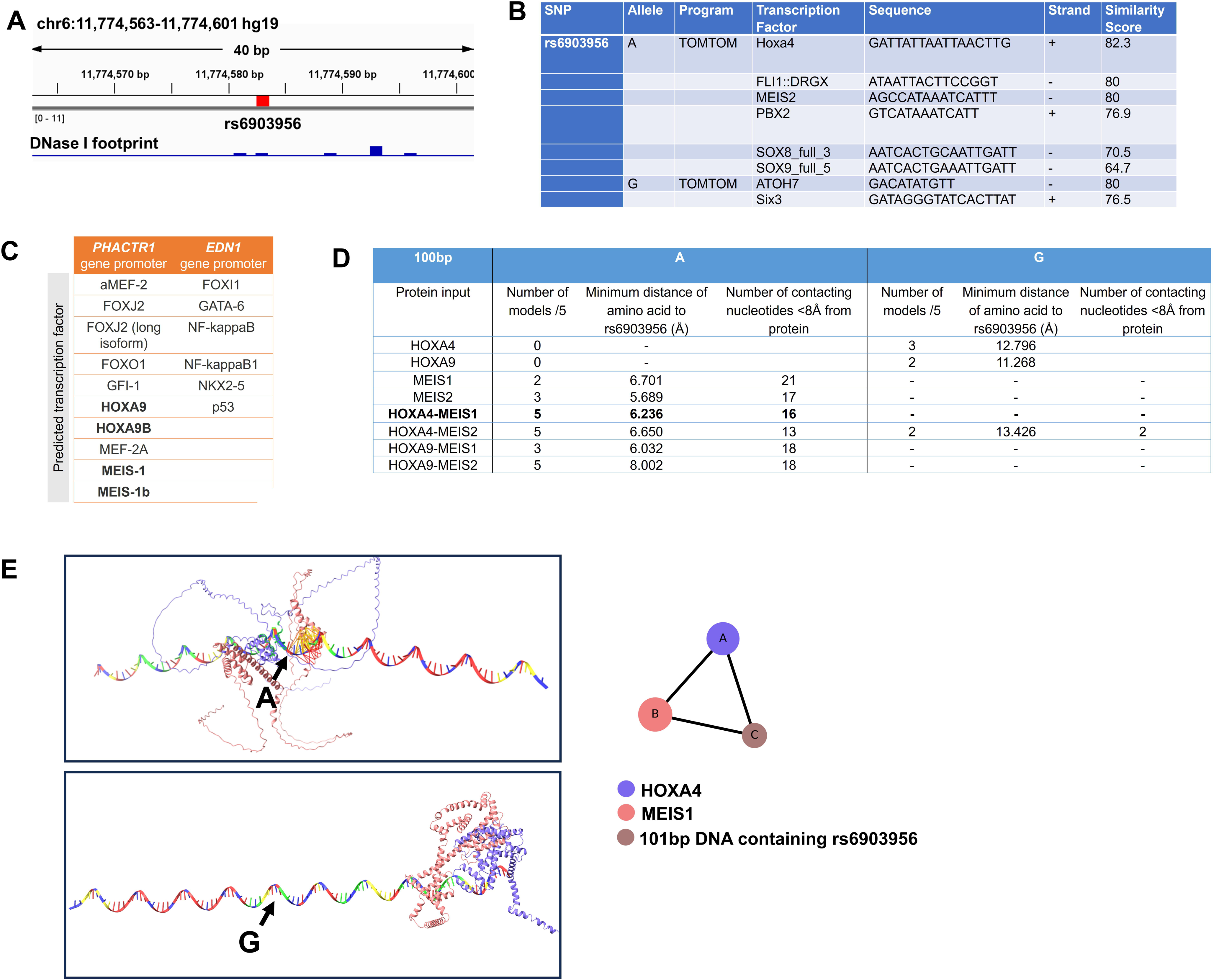
Transcription Factor Motif and Structural Predictions for rs6903956 and HOX/MEIS Proteins. (A) DNase I footprinting analysis (ENCODE Project, GEO accession: GSM1014528) of human umbilical vein endothelial cells on the hg19 chr6:11,774,470-11,774,700 region, highlighting potential binding sites near rs6903956. (B) Predicted transcription factor motifs surrounding rs6903956 based on a 31-bp sequence, analyzed using TOMTOM platform. The similarity score reflects the degree of alignment between the motif and the input sequence. (C) Predicted binding of transcription factors to the *PHACTR1* and *EDN1* gene promoters, as determined using the GeneCards Enhancer Database and QIAGEN TF binding site prediction tools. (D) Structural predictions of the rs6903956 A/G variant and flanking sequences, with HOXA4, HOXA9, MEIS1, and MEIS2 transcription factors and their complexes. AlphaFold 3 was used to predict interactions with a 101-bp region flanking rs6903956 (with the variant at the 50^th^ nucleotide). The output shows results from five AlphaFold models, with the number of models supporting significant binding (<8 Å minimum distance from rs6903956) indicated. Nucleotides within <8 Å of the transcription factor are considered part of the binding site. (E) Structural model of HOXA4 and MEIS1 binding to the 101-bp sequence flanking rs6903956 A (top) and G (bottom). HOXA4 is represented as blue structure, MEIS1 as red structure and the 101-bp DNA strand containing rs6903956 in rainbow colour. The triangle structure illustrates the interaction of the ternary complex of HOXA4, MEIS1 and the 101-bp DNA strand.

Next, we employed the TOMTOM algorithm for *in silico* TF binding site prediction^43^. The risk ‘A’ allele at rs6903956 was predicted to enhance binding affinity for TFs such as HOXA4, MEIS2, PBX2, and SOX proteins (**Fig. 2B**). HOXA4 and MEIS2 are particularly intriguing candidates due to their roles in cardiovascular biology. HOX proteins are well-known regulators of developmental processes and cardiovascular diseases^44^. While HOX proteins typically exhibit low-affinity DNA binding independently, they achieve high-affinity and stable interactions when paired with TALE homeodomain proteins such as MEIS2, which enhance cooperative binding and regulatory function^45^. Interestingly, HOXA4 and MEIS2 were predicted to bind concurrently to opposite strands of the motif containing the ‘A’ risk allele (**Fig. 2B**), highlighting the potential for cooperative regulatory activity or cooperative HOX-MEIS binding^46^. To explore how the localization of rs6903956 within a DHS and its preferential binding to TFs such as HOXA4 and MEIS2 might modulate long-range gene expression, we analyzed TF binding sites in the promoter regions of *EDN1* and *PHACTR1* (**Fig. 2C**). Using the GeneCards Enhancer Database and QIAGEN TF binding site prediction tools, we identified binding motifs for HOXA9 and MEIS1 in the *PHACTR1* promoter. In contrast, no binding motifs for similar TFs were identified in the *EDN1* promoter. This suggests that rs6903956 likely does not regulate *EDN1* expression through HOX-MEIS-mediated mechanisms. Instead, the findings may point to a specific regulatory pathway for *PHACTR1* that involves cooperative binding of HOX and MEIS proteins.

To investigate the molecular mechanisms underlying TF-DNA interactions at the rs6903956 locus, we utilized AlphaFold 3 ^22^, an advanced deep-learning algorithm for 3D molecular interaction modeling. A 101-base pair DNA sequence centered on rs6903956 (position 50) was analyzed for potential interactions with specific HOX (HOXA4, HOXA9) and MEIS (MEIS1, MEIS2) proteins. Simulations were performed for both ‘G’ and ‘A’ alleles at rs6903956, generating five plausible structural models for each condition to explore diverse conformational possibilities. To assess the likelihood of physical or functional interactions, we measured the minimum spatial distance between rs6903956 and the nearest amino acid residue of the interacting TFs, with distances below 8 Å defined as “in contact” ^22^. AlphaFold 3 simulations revealed distinct binding affinities based on the rs6903956 alleles. For the ‘G’ allele, HOXA4 and HOXA9 exhibited low-affinity binding, with minimum distances of 12.796 Å and 11.268 Å, respectively (**Fig. 2D**). In contrast, the ‘A’ allele demonstrated high-affinity interactions with MEIS1 and MEIS2, showing minimum distances of 6.701 Å and 5.689 Å, respectively.

Interestingly, when modeled as cooperative TF complexes, HOX and MEIS proteins formed ternary complexes with preferential binding specificity for the rs6903956 ‘A’ allele (**Fig. 2D**). Notably, HOXA4 and HOXA9, which initially showed weak or no binding to rs6903956 alone, transitioned to high-affinity binding in the presence of MEIS1 or MEIS2. This suggests that MEIS proteins facilitate the recruitment and stabilization of HOX proteins at the rs6903956 A allele binding site. Among the TF complexes, the HOXA4-MEIS1-101bp rs6903956 A DNA complex demonstrated the highest binding affinity, with a minimum distance of 6.236 Å between rs6903956 and the nearest interacting amino acid, indicating this configuration as the most stable and specific. Further structural analysis revealed a robust ternary complex formation between HOXA4, MEIS1, and rs6903956 ‘A’ DNA (**Fig. 2E**, top). This direct interaction among all three components underscores the functional specificity of the ‘A’ allele in facilitating cooperative TF binding. In contrast, the rs6903956 ‘G’ allele DNA failed to support binding of HOXA4 and MEIS1 in a comparable manner (**Fig. 2E**, bottom). These structural predictions highlight the pivotal role of the rs6903956 ‘A’ allele in modulating transcriptional regulation by enabling cooperative binding of HOXA4 and MEIS1.

### Enhancer role of rs6903956 ‘A’ allele in HOXA4-MEIS1-dependent regulation of PHACTR1 in endothelial cells

This enhanced binding may drive allele-specific regulatory effects, as the rs6903956 ‘A’ allele creates novel binding sites for HOXA4 and MEIS1, key regulators of cardiovascular development and endothelial function. To experimentally validate the binding of HOXA4 and MEIS1 to the rs6903956 ‘A’ allele *in situ*, we performed chromatin immunoprecipitation (ChIP) assays using specific antibodies against these transcription factors in WT (AA), UNΔ (AA), and Δ (GG) arterial endothelial cells (**Fig. 3A**). The immunoprecipitated chromatin was quantified using TaqMan qPCR, targeting the genomic region containing rs6903956 (**Supplemental Fig. 2A, B**). ChIP-TaqMan qPCR results revealed significantly stronger binding of HOXA4 and MEIS1 to the rs6903956 ‘A’ allele in WT (AA) and UNΔ (AA) ECs compared to the ‘G’ allele in Δ (GG) ECs (**Fig. 3B**). These results confirm that the rs6903956 ‘A’ allele creates a transcription factor binding site, enabling allele-specific interaction with HOXA4 and MEIS1.

**Figure 3.**
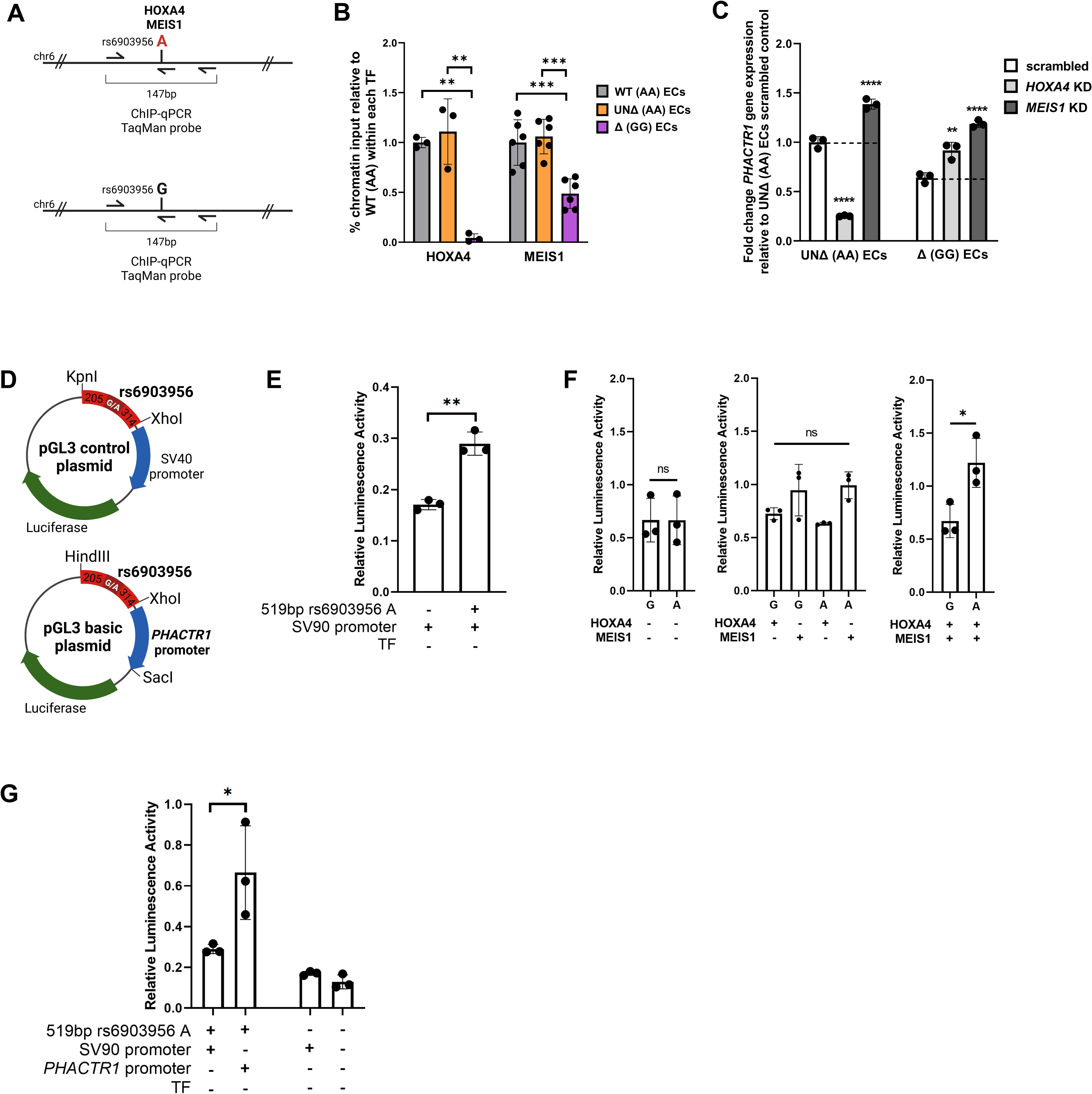
ChIP-TaqMan qPCR, siRNA Knockdown, and Dual Luciferase Assays to Validate the Enhancer Role of rs6903956. (A) Schematic of the ChIP-qPCR assay design targeting the proposed rs6903956 enhancer region. Chromatin immunoprecipitation (ChIP) was performed on wild-type (WT) AA, unedited (UNΔ) AA, and edited (Δ) GG endothelial cells using anti-HOXA9 and anti-MEIS1/2 antibodies. TaqMan qPCR was conducted to amplify a 147-bp region flanking rs6903956. (B) Bar graph showing the percentage of chromatin input for HOXA9 and MEIS1 binding in WT AA, UNΔ AA, and Δ GG ECs, relative to WT AA aECs. Data are presented as means ± S.D. (1-2 donor cell lines, n=3 technical replicates/ donor). One-way ANOVA with post-hoc Tukey’s tests determines statistical significance among three experimental groups within each TF, \*\**p* < 0.01, \*\*\**p* < 0.001. (C) Quantitative RT-PCR analysis of *PHACTR1* expression following siRNA-mediated knockdown of HOXA4 and MEIS1 in UNΔ (AA) ECs and Δ (GG) ECs. Expression levels were normalized to its own scrambled siRNA controls. Bar graphs show means ± S.D. (n = 3 biological replicates). One-way ANOVA with post-hoc Tukey’s tests determines statistical significance relative to its own scrambled, \*\**p* < 0.01, \*\*\*\**p* < 0.0001. (D) Schematic of the pGL3 plasmid constructs used in the dual-luciferase assays. The pGL3 control plasmid contains a 519-bp region flanking rs6903956 cloned under the SV40 promoter, while the pGL3 basic plasmid includes the 519-bp region and the PHACTR1 promoter. These constructs were employed to assess enhancer activity. (E) Relative luciferase activity of pGL3 control plasmids, with and without the 519-bp region flanking rs6903956 ‘A’, transfected into human umbilical vein endothelial cells. The effect of the enhancer region on firefly luciferase activity, normalized to Renilla luciferase activity (internal control), is shown. Data are presented as means ± S.D. (n = 3 biological replicates). Statistical significance between the two groups was determined by two-tailed t-test, \*\**p* < 0.01. (F) Relative luciferase activity of pGL3 basic plasmids containing the 519-bp region flanking rs6903956 (A and G alleles) and the *PHACTR1* promoter. The plasmids were transfected into human umbilical vein endothelial cells with single or co-transfections of HOXA4 and MEIS1. The far-right bar represents the pGL3 basic plasmid control without the enhancer or promoter. Firefly luciferase activity was normalized to Renilla luciferase activity. Data are presented as means ± S.D. (n = 3 biological replicates). Statistical significance between the two groups was determined by two-tailed t-test and one-way anova with post hoc tukey test, \**p* < 0.05; ns, non-significant. (G) Relative luciferase activity of pGL3 control plasmids with the 519-bp region flanking rs6903956 ‘A’ cloned under SV40 or *PHACTR1* promoter control. Firefly luciferase activity was normalized to Renilla luciferase activity. Data are presented as means ± S.D. (n = 3 biological replicates). Statistical significance was determined by two-tailed t-tests, *p ≤ 0.05.

To test the hypothesis that the rs6903956 ‘A’ allele upregulates *PHACTR1* via HOXA4 and MEIS1 binding, we performed siRNA-mediated knockdowns of *HOXA4* and *MEIS1* in UNΔ (AA) and Δ (GG) endothelial cells (ECs) (**Supplemental Fig. 2C**) and measured *PHACTR1* expression by qRT-PCR. In UNΔ (AA) ECs, *HOXA4* knockdown significantly reduced *PHACTR1* expression compared to the scrambled control (**Fig. 3C**), whereas no reduction was observed in Δ (GG) ECs. This confirms that *HOXA4*-mediated regulation of *PHACTR1* is specific to the rs6903956 ‘A’ allele. Despite ChIP-qPCR confirming MEIS1 binding to the rs6903956 A allele, *MEIS1* knockdown unexpectedly led to a significant upregulation of *PHACTR1*. This suggests that MEIS1 may play a modulatory rather than a direct transcriptional role in regulating *PHACTR1*. MEIS proteins are often cofactors that stabilize HOX-DNA interactions rather than independently driving transcription^47^. Functional redundancy within the MEIS family, such as compensatory effects from MEIS2, may also account for this observation. Collectively, these results support a model in which the G-to-A substitution at rs6903956 creates a TF binding site for HOXA4. In Δ (GG) ECs, the absence of this TF binding site prevents the engagement of this regulatory pathway.

To validate the regulatory role of rs6903956 in facilitating enhancer-promoter interactions that drive *PHACTR1* expression, we performed a dual-luciferase assay using a 519 bp DNA fragment flanking rs6903956. This fragment was cloned into a pGL3-Control vector upstream of an SV40 promoter and transfected into human primary endothelial cells (**Fig. 3D**, top; **Supplemental Fig. 2D**, top). The rs6903956 ‘A’ allele fragment significantly increased luciferase activity compared to the control plasmid lacking the enhancer region, confirming that the rs6903956 ‘A’ allele fragment confers enhancer activity to the SV40 promoter (**Fig. 3E**).

To further investigate the allele-specific regulation of the *PHACTR1* promoter and the cooperative roles of HOXA4 and MEIS1, the same 519 bp fragment (A/G alleles) was cloned upstream of the *PHACTR1* promoter in the pGL3-Basic vector (**Fig. 3D**, bottom; **Supplemental Fig. 2D**, bottom). Co-transfections were performed in human primary endothelial cells using HOXA4 and MEIS1 expression plasmids, individually and in combination. In the absence of transcription factors, there was no significant difference in luciferase activity between enhancer regions flanking the rs6903956 ‘A’ or ‘G’ alleles (**Fig. 3F**), indicating that the enhancer activity is transcription factor dependent. Expression of HOXA4 alone did not significantly enhance transcriptional activity, consistent with AlphaFold predictions that HOXA4 requires MEIS proteins for high-affinity binding to the rs6903956 ‘A’ allele. Similarly, MEIS1 expression alone resulted in a modest but non-significant increase in transcriptional activity for both alleles. However, co-expression of HOXA4 and MEIS1 significantly increased transcriptional activity for the enhancer region flanking the rs6903956 ‘A’ allele compared to the ‘G’ allele (**Fig. 3F**), highlighting the critical role of the HOXA4-MEIS1 complex in activating the *PHACTR1* promoter in an allelespecific manner.

Notably, transcriptional activity driven by the enhancer region flanking rs6903956 ‘A’ allele with the *PHACTR1* promoter was significantly higher than that observed with the SV40 promoter (**Fig. 3G**), suggesting that rs6903956 functions as a locus-specific enhancer regulating *PHACTR1* expression in endothelial cells. In summary, these findings establish that the rs6903956 ‘A’ allele region functions as an enhancer that facilitates HOXA4 and MEIS1 recruitment, driving *PHACTR1* expression in endothelial cells.

### Allelic variation at rs6903956 drives differential endothelial activation and monocyte adhesion

To directly link genetic variation to endothelial phenotypes, we investigated the functional consequences of rs6903956 relating to its association with *PHACTR1*. Endothelial *PHACTR1* has been shown to mediate inflammation by activating NF-κB-dependent *ICAM-1* expression under disturbed flow, with silencing of *PHACTR1* resulting in reduction of *ICAM-1* levels in human endothelial cells^35^. Conversely, *PHACTR1* overexpression promotes inflammation and monocyte adhesion^48^. Based on this, we postulated that UNΔ (AA) ECs, expressing higher *PHACTR1* levels compared to Δ (GG) ECs, would exhibit enhanced inflammatory activation. In static cell culture condition, UNΔ (AA) ECs exhibited a more pronounced pro-inflammatory phenotype, characterized by higher mean fluorescence intensities of ICAM-1 compared to Δ (GG) ECs (**Fig. 4A**). This observation suggests that the rs6903956 risk allele may drive heightened endothelial activation and inflammation, consistent with our previously reported association of the risk allele ‘A’ with elevated endothelial injury biomarkers in CAD patients ^11^.

**Figure 4.**
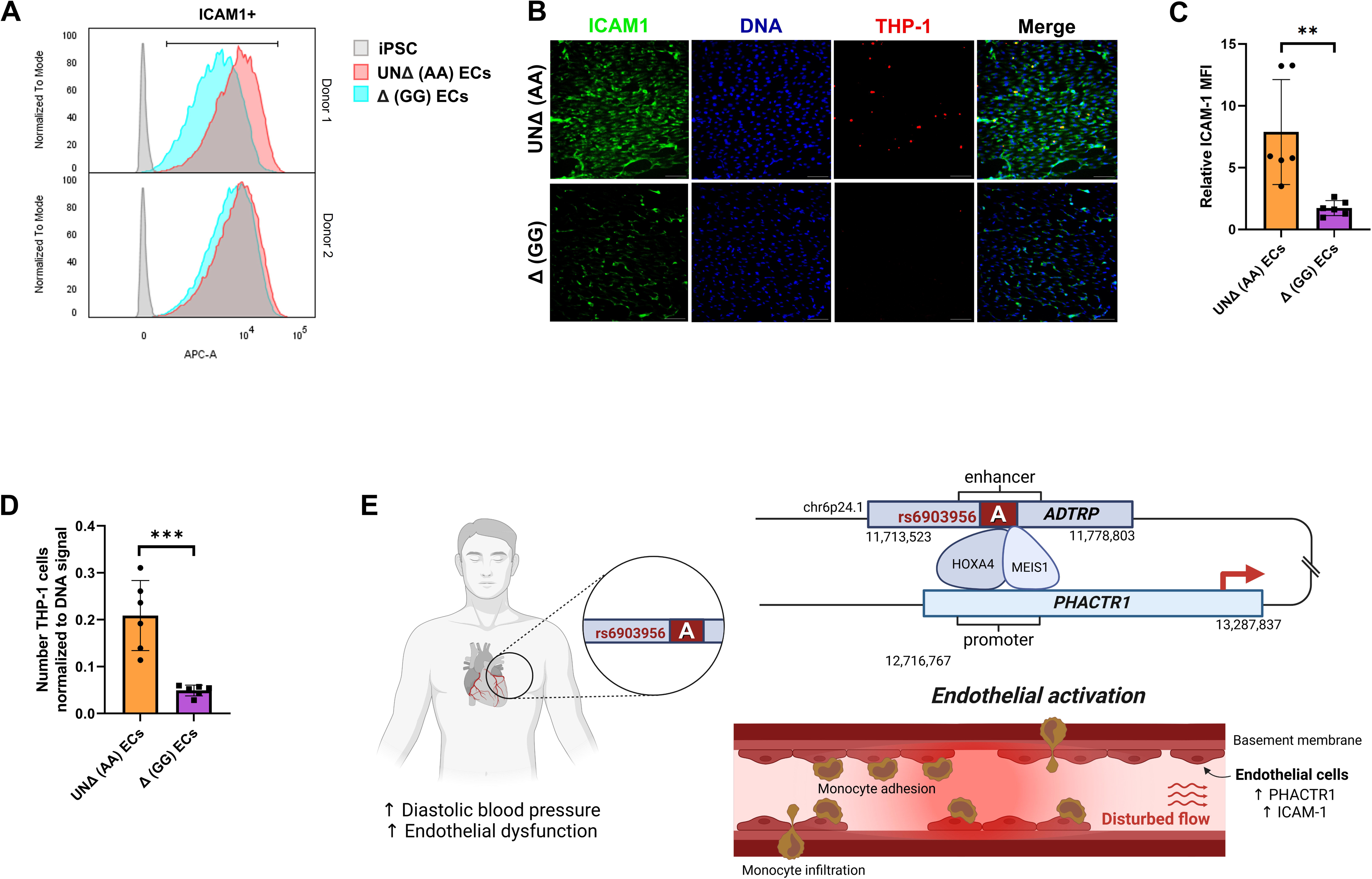
ICAM1 Expression and Monocyte Adhesion in Co-Culture with **Δ** (GG) Compared to UN**Δ** (AA) Endothelial Cells. (A) Flow cytometry analysis of proinflammatory marker ICAM1 expression on unedited UNΔ (AA) ECs and edited Δ (GG) ECs, highlighting differential expression profiles (mean fluorescence intensity) between the two genotypes across two donor samples. (B) Representative immunofluorescence images of ICAM1 (green), DAPI (blue), and adhered THP-1 monocytes (red) on UNΔ (AA) and Δ (GG) endothelial cells grown under disturbed flow condition. Scale bar: 100 μm. (C) Quantification of ICAM1 expression in UNΔ (AA) ECs and Δ (GG) ECs, represented as mean fluorescence intensity (MFI). Bar graphs represent means ± S.D. (n = 3 biological replicates/ donor from 2 donors). Mann-Whitney U test was performed to compared between two cell lines, \*\**p* < 0.01. (D) Quantification of THP-1 monocyte adhesion on UNΔ (AA) ECs and Δ (GG) ECs, represented as number of THP-1 cells normalized to DNA signal. Bar graphs represent means ± S.D. (n = 3 biological replicates/ donor from 2 donors). Two-tailed t-tests was performed to compared between two cell lines, \*\*\**p* ≤ 0.001. (E) Mechanistic model.

Under disturbed flow, UNΔ (AA) and Δ (GG) ECs were exposed to non-uniform flow with low shear stress (±6 dynes/cm², 1 Hz) in the presence of THP-1 monocytes. Immunofluorescence staining for ICAM-1 and quantification of adhered monocytes were performed (**Fig. 4B**). UNΔ (AA) ECs displayed significantly higher ICAM-1 expression and greater extent of monocyte adhesion compared to Δ (GG) ECs (**Fig. 4C, D**). These findings demonstrate an allele-specific endothelial phenotypic response to disturbed flow, with UNΔ (AA) ECs exhibiting elevated endothelial activation hallmarks relative to Δ (GG) ECs (**Fig. 4E**).

## Discussion

GWAS have identified thousands of non-coding variants associated with human disease risk, but understanding their regulatory mechanisms remains a major challenge. Approaches like PheWAS, eQTL mapping, and genome annotation tools have helped elucidate the functions of risk variants, but they often lack cell-type specificity and require experimental validation to confirm causal relationships. In this study, we show that rs6903956 influences *PHACTR1* expression via a cooperative interaction between HOX and MEIS proteins. The G-to-A substitution at rs6903956 enhances transcription factor complex binding with the ‘A’ alleles, facilitating longrange enhancer-promoter interactions that activate *PHACTR1*. Our approach combined population-scale data, genome editing, and functional assays in a human endothelial cell model to establish a link between rs6903956 and endothelial dysfunction relevant to CAD pathogenesis.

The rs6903956 variant was identified as a susceptibility locus for CAD in the Chinese Han population, with the ‘A’ risk allele associated with decreased *ADTRP* expression in peripheral blood leukocytes^6^. Luo et al. then proposed that the ‘A’ allele disrupts the GATA2 binding site within an enhancer region, reducing *ADTRP* promoter activity^8^. However, in our base-edited arterial endothelial cell model, rs6903956 did not significantly affect *ADTRP* expression, suggesting cell-typespecific effects of the variant. This discrepancy highlights the complexity of interpreting rs6903956’s role across different cell types involved in CAD pathogenesis, such as macrophages, endothelial cells and smooth muscle cells.

Although we did not observe a direct link between rs6903956 and *ADTRP* expression in endothelial cells, our findings suggest that the rs6903956 ‘A’ allele influences endothelial function through *ADTRP*-independent mechanisms, highlighting the complexity of the 6p24.1 locus in CAD. Notably, the ‘A’ allele was associated with increased expression of *PHACTR1*, a gene located approximately 942 kb downstream of rs6903956 within the same topologically associating domain. *PHACTR1*, an actin-and protein phosphatase 1 (PP1)-binding protein, is implicated in F-actin remodeling and PP1 regulation, with its locus at 6p24.1 linked to susceptibility to several vascular diseases, including CAD^40,49–54^. Initially identified in endothelial cells via suppression subtractive hybridization, *PHACTR1* has been shown to play a critical role in endothelial function. Knockdown of *PHACTR1* in endothelial cells impairs tube formation and induces apoptosis, as well as reduces expression of adhesion molecules such as ICAM-1 and VCAM-1^55,56^, underscoring its role in oxidative stress and inflammation. Furthermore, *PHACTR1* contributes to disturbed flow-mediated atherosclerosis by repressing PPARγ activity^35^.

The role of *PHACTR1* in CAD is complex and multifaceted, with evidence suggesting both proatherogenic and potential atheroprotective effects^57^. The lead SNP on *PHACTR1*, rs9349379, has been linked to higher *EDN1* protein levels^36^, which encodes endothelin-1, a potent vasoconstrictor peptide produced by endothelial cells. Overexpression of *PHACTR1* has been shown to upregulate *EDN1* and downregulate eNOS expression in human endothelial cells, reducing NO production^48^. While the precise mechanism behind *PHACTR1*-mediated *EDN1* upregulation remains unclear, we observed a similar increase in *EDN1* levels in our genetically corrected endothelial cells with GG non-risk genotype, suggesting that the rs6903956 ‘A’ allele, which drives *PHACTR1* upregulation, may contribute to proinflammatory and vasoconstrictive pathways by elevating *EDN1*. Our findings, combined with the regulatory effects of known *PHACTR1* SNPs, highlight the intricate and dynamic regulatory landscape at chromosome 6p24.1.

Our study underscores the pivotal role of the rs6903956 ’A’ allele in modulating transcriptional regulation through cooperative binding of HOXA4 and MEIS1. *In silico* transcription factor analyzes, based on Position Weight Matrices (PWMs), quantify binding likelihood across sequences but often face challenges in identifying specific proteins within similar families due to structural overlap. To refine these predictions, we utilized AlphaFold 3 to model protein-DNA interactions at rs6903956, focusing on amino acid proximity and structural stability. Among potential complexes, HOXA4-MEIS1 emerged as the most likely interactor with the rs6903956 enhancer. This is supported by ChIP-qPCR detection of MEIS1 binding to the enhancer, although MEIS1 knockdown alone did not significantly affect *PHACTR1* expression, suggesting a nuanced regulatory role. Notably, a HOXA9-PBX2-MEIS1 complex was previously reported in myeloid leukemia cells, where MEIS1 inclusion did not enhance transcriptional activity^58^, consistent with our findings. Furthermore, we acknowledge potential compensatory roles of other HOX and TALE family members, such as PBX2 and MEIS2, as functional redundancy is well-documented among HOX paralogues^59^. These observations suggest that rs6903956-mediated transcriptional regulation may involve multiple protein complexes, highlighting the intricate regulatory dynamics of this enhancer region.

Our investigation of the 6p24.1 risk locus, identified by GWAS as a CAD susceptibility locus, has uncovered critical cis-regulatory mechanisms that drive endothelial cell dysfunction. Using base editing and arterial endothelial cell models, we demonstrated that the rs6903956 ‘A’ allele functions as an enhancer, promoting *PHACTR1* expression via cooperative interactions with HOX and TALE family transcription factors. These findings illuminate the intricate genetic architecture of the 6p24.1 locus and provide novel insights into transcription factor dynamics in cardiovascular disease, connecting rs6903956 to pathways contributing to vascular inflammation. By bridging functional genetics with disease-relevant cell models and molecular underpinnings of 6p24.1, our study enhances the standard for interrogating GWAS variants, advancing genetic testing and precision medicine.

## Acknowledgements

We thank all patients and healthy donors who have participated in this study. Special thanks to the team at National University Hospital for coordinating clinical sample collection. The HELIOS study is also supported by an outstanding team of administrative and operational staff.

## Sources of Funding

C.C. was funded by an Academic Research Fund (2018-T1-001-030, MOE-T2EP30122-0018) from the Ministry of Education, Singapore, and Vascular Research Initiative from Lee Kong Chian School of Medicine, Singapore. K.Y.T. is supported by NTU Research Scholarship. J.L. and M.L are supported by Singapore Ministry of Health’s (MOH) National Medical Research Council (NMRC) under its OF-LCG funding scheme (MOH-000271-00), Singapore Translational Research (StaR) funding scheme (NMRC/StaR/0028/2017), the National Research Foundation, Singapore through the Singapore MOH NMRC and the Precision Health Research, Singapore (PRECISE) under the National Precision Medicine programme (NMRC/PRECISE/2020) and intramural funding from Nanyang Technological University, Lee Kong Chian School of Medicine and the National Healthcare Group. RNA sequencing was partially funded by i) Ministry of Education Academic Research Fund Tier 1 Grant (RS09/20), ii) A*STAR-NHMRC Joint Grant Call (A20PRb0138), iii) Start-Up Grant (awarded to M.L.) from Lee Kong Chian School of Medicine, Nanyang Technological University, Singapore and iv) Imperial - Nanyang Technological University Collaboration Fund (awarded to M.L.).

MESA dataset was applied for through dbGAP via dbGaP study accession phs000209.v13.p3 and phs000420.v6.p3. MESA and the MESA SHARe project are conducted and supported by the National Heart, Lung, and Blood Institute (NHLBI) in collaboration with MESA investigators. Support for MESA is provided by contracts N01-HC95159, N01-HC-95160, N01-HC-95161, N01-HC-95162, N01-HC-95163, N01-HC-95164, N01-HC-95165, N01-HC95166, N01-HC-95167, N01-HC-95168, N01-HC-95169, UL1-RR-025005, and UL1-TR-000040. Funding for SHARe genotyping was provided by NHLBI Contract N02-HL-64278. Genotyping was performed at Affymetrix (Santa Clara, California, USA) and the Broad Institute of Harvard and MIT (Boston, Massachusetts, USA) using the Affymetrix Genome-Wide Human SNP Array 6.0. Funding support for the FMD dataset was provided by grant and contract numbers N01 HC-95159-69, RR-024156, and HL077612. The China cohort was supported by the National Natural Science Foundation of China (82304098) and the Major Science and Technology Projects of Guangxi, China (AA22096026).

## Competing interests

The authors declare no competing interests.

## Data Availability Statement

The authors declare that all data supporting the findings of this study are available within the paper and supplementary information.

## Authors’ contributions

K.Y.T. contributed to conceptualization, methodology, formal analysis, investigation, writing of original draft, and visualization. H.S.W., N.N., V.K.W. and K.L.L. contributed to methodology, formal analysis, and visualization. M.I.A. and R.S.Y.F. contributed to methodology and resources. D.T., Y.W., X.G., J.L., M.L. contributed to formal analysis and data curation. C.K.H. and M.Y.Y.C. contributed to resources and data curation. C.C. contributed to project administration, conceptualization, formal analysis, investigation, resources, visualization, writing original draft, supervision, and funding acquisition. All authors were involved in the review and editing of the manuscript.

## Declaration of generative AI in the writing process

During the preparation of this work the authors used ChatGPT in order to improve language and readability. After using this tool, the authors reviewed and edited the content as needed and take full responsibility for the content of the publication.

## Notes

### Competing Interest Statement

The authors have declared no competing interest.

