## Supplemental for "Coronary Artery Disease Risk Variant rs6903956 Links to Endothelial Dysfunction via *PHACTR1* Regulation"

[1]Lee Kong Chian School of Medicine, Nanyang Technological University, Singapore.
[2]Yong Loo Lin School of Medicine, National University of Singapore, Singapore
[3]Genome Institute of Singapore, Agency for Science, Technology and Research, Singapore.
[4]Department of Occupational and Environmental Health Sciences, School of Public Health, Peking University, Beijing, China
[5]Khoo Teck Puat, National University Children's Medical Institute, National University Health System, Singapore
[6]National University Heart Centre, National University Health System, Singapore
[7]North Region, Institute of Mental Health, Singapore
[8]Department of Epidemiology and Biostatistics, Imperial College London, London, UK
[9]National Skin Centre, Singapore
[10]Institute of Molecular and Cell Biology, Agency for Science Technology and Research, Singapore.

### Supplementary Figures

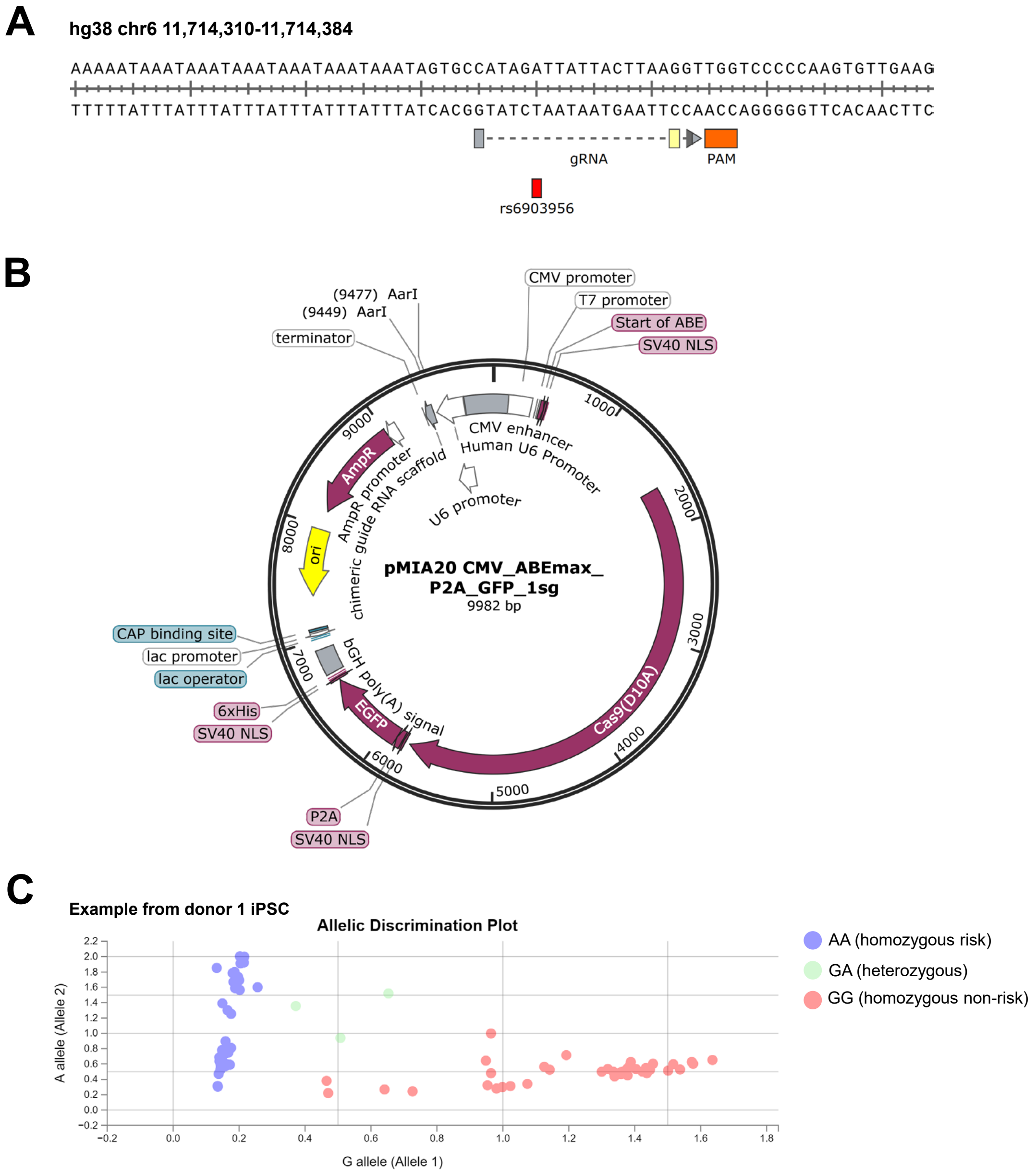


**Supplemental Figure 1. pMIA20 Adenosine Base Editing Plasmid Design and Validation of Base-Edited Clones.** (**A**) Design of guide RNA targeting rs6903956 for base editing, highlighting the NGG PAM site and the rs6903956 variant in red. (**B**) pMIA20 plasmid schematic, featuring an empty backbone that expresses the sgRNA cassette (AarI cloning site), a human U6 promoter-driven D10A Cas9, and EGFP for detection of successful transfection. This plasmid will be made available in the Addgene. (**C**) Allelic discrimination plot from a rs6903956 TaqMan genotyping assay, where each dot represents a cell line expanded from a single FACS-isolated cell. Samples containing the A allele (homozygous) are colored blue, G allele (homozygous) samples are red, and heterozygous (GA) samples are green.

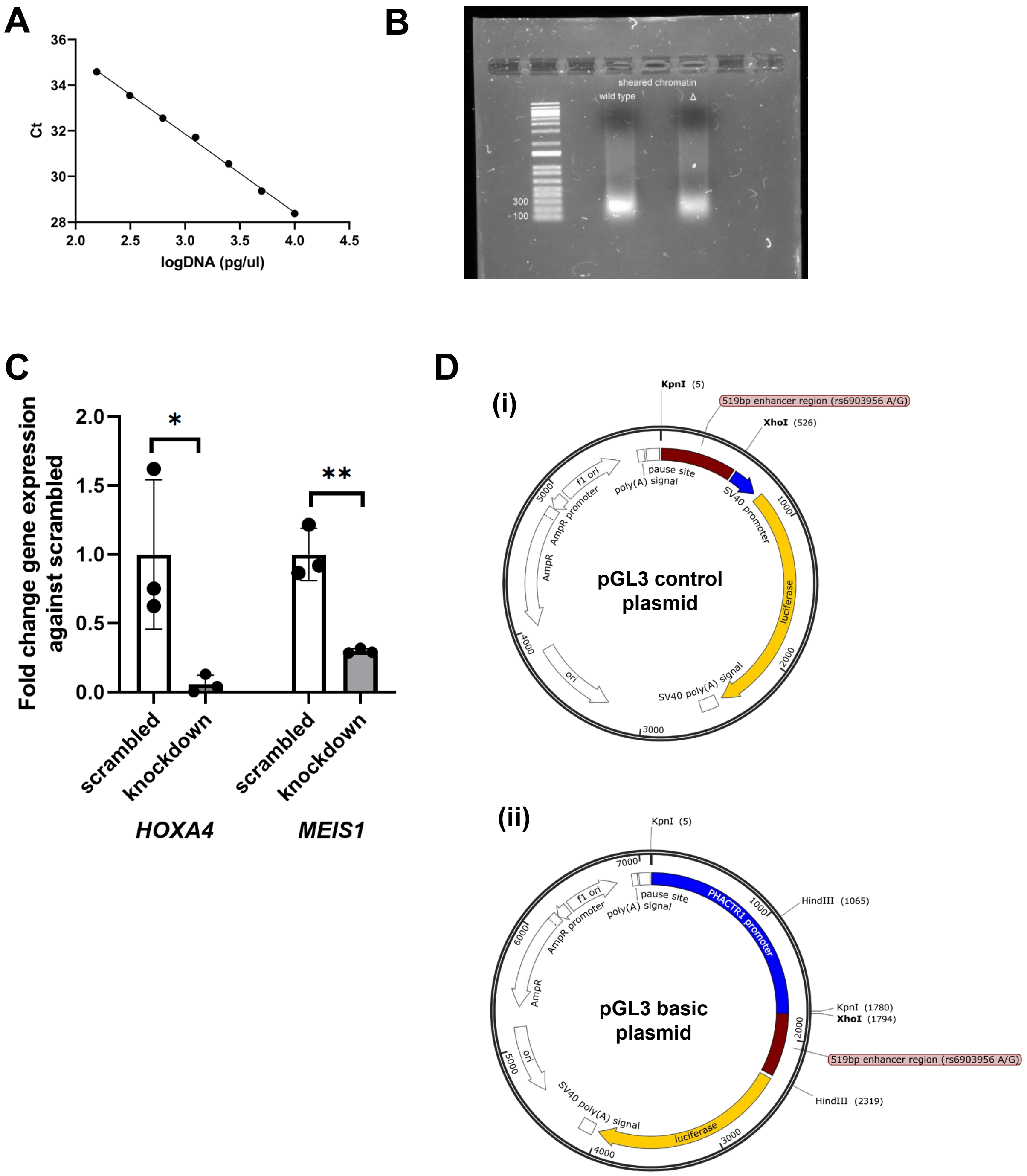


**Supplemental Figure 2. Validation of ChIP-TaqMan qPCR, siRNA Knockdown, and Schematic Representation of pGL3 Plasmids.** (**A**) Primer dilution curve for ChIP-TaqMan qPCR primers, demonstrating specificity and efficiency of the primers. The analysis was performed using genomic DNA from iPSC donor 1. (**B**) Gel electrophoresis image showing chromatin DNA sheared to an estimated size of 100–300 bp through formaldehyde fixation and probe sonication. This sheared DNA was used for ChIP-TaqMan qPCR. (**C**) Fold-change in gene expression of *HOXA4* and *MEIS1* relative to scrambled siRNA control, measured using qRT-PCR at 72 hours post-siRNA knockdown using Dharmacon's pooled siRNA at 5 nM concentration. Expression levels were normalized to the scrambled control. Bar graphs represent mean ± S.D. (n = 3 biological replicates). two-tailed t-test was performed to compared between scrambled control and knockdown sample, *$p < 0.05$, **$p < 0.01$. (**D**) Schematic plasmid maps for (i) the pGL3 control plasmid containing the 519-bp fragment flanking rs6903956 (A/G alleles) and (ii) the pGL3 basic plasmid with the 519-bp rs6903956 A/G fragment and the *PHACTR1* promoter inserted. The cloning strategies for these plasmids support dual-luciferase assay validation of enhancer activity.

#### Supplementary Tables

**Supplemental Table 1. Phenome-Wide Association Study (PheWAS) of rs6903956 in Circulatory System Disorders from the UK Biobank.** This table presents the associations between rs6903956 and clinically diagnosed circulatory system disorders, derived from the UK Biobank dataset. Effect sizes are reported alongside their corresponding standard errors (SE). Sample sizes for each diagnosis are indicated as the number of cases and controls analyzed.

| Category | Clinical Diagnosis | *p*-value | Effect size (SE) | Number of samples |
|---|---|---|---|---|
| Circulatory system | **Essential hypertension** | **0.023** | 0.015 (0.0066) | 77465/328796 |
| Circulatory system | **Hypertension** | **0.025** | 0.015 (0.0066) | 77714/328796 |
| Circulatory system | Coronary atherosclerosis | 0.28 | 0.012 (0.011) | 20454 / 374655 |
| Circulatory system | Disease of capillaries | 0.05 | 0.23 (0.12) | 144/3987056 |
| Circulatory system | Nonrheumatic tricuspid valve disorders | 0.073 | 0.25 (0.14) | 105/400662 |
| Circulatory system | Varicose veins of lower extremity, symptomatic | 0.087 | 0.10 (0.058) | 623/367909 |

**Supplemental Table 2. Genotype Information for Known PHACTR1 Risk Variants in our CAD Patients.** This table summarizes the genotypes of our CAD patient donor 1 and donor 2 for previously identified PHACTR1 risk variants associated with blood pressure phenotypes. The genotypic data for each risk variant are provided for both donors.

| ***PHACTR1* SNP Genotyping in our iPSC lines** | | | | | | | |
|---|---|---|---|---|---|---|---|
| | Donor 1 | Donor 2 | Major allele | Minor allele | Disease link | References | GWAS n number |
| Rs9369640 | A | A | A | C | Minor allele C is protective for hypertension | PMC4251179 | n = 5460 |
| Rs1223397 | G | G | G | C | rs1223397 C risk allele is significantly associated with pulse pressure. | PMID:22763476 | Genome-wide linkage and association scans: 63 middle-aged dizygotic twin pairs |
| Rs9349379 | A | A | G | A | In Chinese Han, minor A allele is associated with higher pulse pressure. | PMC7276519 | n=706 |
| Rs9349379 | A | A | A | G | In European: Minor allele G at rs9349379 links to increased vascular disease risk due to higher ET-1 levels. | PMC5785707 | |

**Supplementary Table S3. List of primary antibodies for immunofluorescence and flow cytometry.**

| Antibodies | | | |
|---|---|---|---|
| **Antibodies** | **Manufacturer Cat #, RRID** | **Concentration** | **Application** |
| PECAM1 | Miltenyi Biotec Cat# 130-097-857 | 20 µl ($1X10^7$ cells) | Magnetic beads for sorting iPSC-EC |
| DLL4-APC | BioLegend Cat# 130813 | 1:20 ($1X10^6$ cells) | Flow cytometry for EC marker |
| VECAD (CDH5)-PE | BD Biosciences Cat# 561714, RRID:AB_10895800 | 1:50 ($1X10^6$ cells) | Flow cytometry for EC marker |
| Anti-HOXA4 | Santa Cruz BiotechnologyCat# sc-515418 | 0.8 µg/well | ChIP-qPCR antibody |
| Anti-MEIS1/2/3 | Santa Cruz BiotechnologyCat# sc-101850 | 0.8 µg/well | ChIP-qPCR antibody |
| Anti-CD144 rabbit polyclonal | Abcam Cat# ab33168 | 0.1-1 µg/mL | Immunofluorescence |
| Goat anti-rabbit IgG, Alexa Fluor™ 488 | Invitrogen Cat# A-11011 | 2 µg/mL | Immunofluorescence |
| ICAM-1 mouse monoclonal | Invitrogen Cat# MA5407 | 1:250 | Immunofluorescence |
| Goat anti-mouse IgG Alexa Fluor™ 488 | Invitrogen, Cat# A-11029 | 1:400 | Immunofluorescence |

**Supplementary Table S4. qPCR Primer sequences.**

| Primers for qPCR and genotyping | | |
|---|---|---|
| **Primer for Gene/ Risk Variants** | **Forward Sequence** | **Reverse Sequence** |
| *GAPDH* | CCGTCAAGGCTGAGAACGG | CTCAGCGCCAGCATCGC |
| *HOXA4* | TGTCAGCGCCGTTAACCC | GCCGGGTCAGGTATCGATTG |
| *MEIS1* | CCAGTCCAACCGAGCAGTAA | CACTCATAGGTCCTGGTGCT |
| *PHACTR1* | CGAAGACGACGACAGCTCAT | TTCTTCCAGCTCTCGCTTGG |
| *EDN1* | AAGGCAACAGACCGTGAAAAT | CGACCTGGTTTGTCTTAGGTG |
| *ADTRP* | GAAGACAGGACTCACCTTGCTG | GGCAAACACAGGATACACCCAG |
| *TaqMan qPCR primer for rs6903956* | GCCTGTGAACAGTGAGAAGA | GGGAACAGAGAGAGATTCCATC |
| *TaqMan qPCR probe for rs6903956* | CCAAGTGTTGAAGTGGGTGCCATACT | - |
| *rs9369640* | GCCTCCACTAGGGACAACAC | CTTTCCCTGCTGCTTCTCCA |
| *rs1223397* | GCATGTGGCACAGTAGGGAT | AAATGGGGAGCCACTGAAGG |
| *rs9349379* | CACCGTTACCGGGACCATTT | CTTGGCGAGTTCACTGCATC |

**Supplementary Table S5. Primers for fragment amplification (plasmid construction)**

| | |
|---|---|
| AarI oligo F | CACCGTTGGCAGGTGGACACCTGCTTCT |
| AarI oligo R | AAACAGAAGCAGGTGTCCACCTGCCAAC |
| 1sg into ABEmax F | gcgatgtacgggccagatatAAAAAAGCACCGACTCGG |
| 1sg into ABEmax R | tagtcaataatcaatgtcaacgcgtGAGGGCCTATTTCCCATG |
| Allele-Specific qPCR F | GGGGACCAACCTTAAGTAATAACCTATG |
| Allele-Specific qPCR R | GAGCCAAGATTGTGCCACTGC |
| TaqMan qPCR F | GCCTGTGAACAGTGAGAAGA |
| TaqMan qPCR R | GGGAACAGAGAGAGATTCCATC |
| TaqMan qPCR probe | CCAAGTGTTGAAGTGGGTGCCATACT |
| Guide RNA | CATAGATTATTACTTAAGGT |
